# Delicate balances in cancer chemotherapy: Modeling immune recruitment and emergence of systemic drug resistance

**DOI:** 10.1101/2019.12.12.874891

**Authors:** Anh Phong Tran, M. Ali Al-Radhawi, Irina Kareva, Junjie Wu, David J. Waxman, Eduardo D. Sontag

## Abstract

Metronomic chemotherapy can drastically enhance immunogenic tumor cell death. However, the responsible mechanisms are still incompletely understood. Here, we develop a mathematical model to elucidate the underlying complex interactions between tumor growth, immune system activation, and therapy-mediated immunogenic cell death. Our model is conceptually simple, yet it provides a surprisingly excellent fit to empirical data obtained from a GL261 mouse glioma model treated with cyclophosphamide on a metronomic schedule. The model includes terms representing immune recruitment as well as the emergence of drug resistance during prolonged metronomic treatments. Strikingly, a fixed set of parameters, not adjusted for individuals nor for drug schedule, excellently recapitulates experimental data across various drug regimens, including treatments administered at intervals ranging from 6 to 12 days. Additionally, the model predicts peak immune activation times, rediscovering experimental data that had not been used in parameter fitting or in model construction. The validated model was then used to make predictions about expected tumor-immune dynamics for novel drug administration schedules. Notably, the validated model suggests that immunostimulatory and immunosuppressive intermediates are responsible for the observed phenomena of resistance and immune cell recruitment, and thus for variation of responses with respect to different schedules of drug administration.

## 1 Introduction

Immune system involvement in cancer progression has been well established, leading to increased efforts to harness the ability of the host immune system to fight off growing tumors [1]. Standard of care chemotherapy regimens typically involve drug administration on a maximum tolerated dose (MTD) schedule [2]. These regimens aim to target most cancer cells at once, but frequently lead to a reduction in tumor burden only in the short term [3] and often give rise to drug resistance [4, 5]. One reason for this longer-term failure is that MTD-based treatment can cause collateral damage to the host immune system, thus diminishing its ability to target the tumor [3]. An optimal treatment regimen should strike a balance between drug-induced tumor cell kill and damage to the immune system, allowing the two modes of cancer cell elimination to complement each other. Indeed, high frequency low dose drug administration, also known as metronomic chemotherapy, can in some cases strike the right balance and induce immunogenic cell death (ICD) in tumor tissue by selecting an appropriate choice of drug, dose, and time interval between treatments [6, 7, 8, 9, 10, 11, 12]. Successful achievement of ICD-based therapeutic outcomes during anticancer therapy is dependent on complex interactions between the drug, the tumor, and the host immune system, the nature of which is still being uncovered [13].

To better understand the mechanisms by which metronomic chemotherapy enables an anti-tumor immune response, it is important to understand how tumors are able to evade immunosurveillance in the first place [1]. An important and well-studied evasion route is through the accumulation of mutations and epigenetic modifications that help avoid immunosurveillance [14, 15]. Our mathematical model will focus on network effects, in contrast to such (epi)genetic changes in tumors.

### 1.1 Interaction dynamics between tumors and the immune system

The proposed model is phenomenological, its components representing the combined effects of a variety of immunostimulatory as well as immunosupressive processes. For background, we next briefly discuss some of these immune-related processes, which involve modifications of the tumor microenvironment (TME). Such modifications include increased acidity resulting from altered nutrient metabolism [16, 17] and altered balance between cytotoxic and regulatory immunity through the recruitment by the tumor of immunosuppressive cells, such as regulatory T cells (Tregs) [18, 19, 20, 21, 22] and myeloid-derived suppressor cells (MDSCs) [23, 24].

Some examples of TME modifications are as follows. Macrophages with an M2 phenotype can produce high levels of TGF-*β*, IL-10, and vascular endothelial growth factor (VEGF), promoting tumor growth [25, 26, 27, 28]. In other cases, tumor-derived factors and gangliosides can alter dendritic cell (DC) phenotype leading to lower levels of CD80, CD86, CD40 and high indoleamine 2,3-dioxygenase expression that contributes to suppression of T cell immunity [29]. Immunosurveillance can also be evaded through production of various immunosuppressive cytokines such as TGF-*β* that play an important role in suppressing macrophages and monocytes [30]. Other factors such as tumor necrosis factor (TNF)-*α*, IL-1, IL-6, colony stimulating factor (CSF)-1, IL-8, IL-10, and type 1 interferons (INFs) can also contribute to cancer growth [31, 32, 33, 34, 35]. Additionally, pro-angiogenic factors such as VEGF can inhibit differentiation of progenitors into DCs [36]. IL-10 and TGF-*β* can also inhibit DC maturation. Ganglioside antigens can also suppress cytotoxic T-cells (CTLs) and dendritic cells (DCs) function [37]. Immunosuppressive enzymes such as IDO, arginase, and inhibitor of nuclear factor kappa-B kinase (IKK)2 may also contribute to tumor progression via direct actions on tumor cell proliferation or through induction of T cell tolerance/suppression [38, 39, 40, 41].

### 1.2 Network effects of chemotherapy interventions

By targeting various components that regulate immune tolerance, cancer chemotherapy drugs, such as mitoxantrone, idarubicin, doxorubicin, and cyclophosphamide can induce immunogenic cancer cell death in addition to their direct cancer cell cytotoxic effects [42, 8, 7, 43]. By using an optimized drug dose and schedule of administration, favorable immune responses can be achieved, including increases in macrophage recruitment and maturation [44], proliferation of NK cells, levels of IFN-*γ* [45], as well as elevated post-apoptotic release of the nuclear chromatin binding protein HMGB1, which can stimulate antigen presentation by DCs, helping CD8^+^ T cell activation [46, 47]. In some cases, type-1 interferon signaling pathways are switched on, leading to host antitumor immunity activation [48, 49]. Immunosuppressive molecules, like CD31, CD46, CD47, are downregulated on dendritic cells by ICD treatment [50]. Molecular chaperones such as HSP90 appear on the tumor cell surface, promoting DC maturation [51]. These optimized drug administration conditions can also lead to transient lymphopenia, which upregulates repair mechanisms and can lead to a vast array of immunostimulatory outcomes, including enhanced T-cell activation, immune recruitment, DC differentiation and maturation, as well as the release of large amounts of chemokines and cytokines [52, 53]. Cytotoxic effects on immunoregulatory cells, such as MDSCs and Tregs, contribute to restoration of anti-tumor immunity by decreasing suppression of T-cells and NK cells [54]. Other factors affecting immunogenicity include changes in MHC-1 molecules and tumor-specific antigens on the tumor cell surface [55], stress-induced expression of NK cell stimulatory ligands, and decreases in NK inhibitory ligands [56].

The ability of drugs to induce anti-tumor immune responses is not sufficient by itself to insure a successful therapeutic response, as the effect on various compartments of the immune system, and thus on overall tumor burden can vary dramatically depending on dose, schedule and tumor type. Scheduling and dosing of an ICD drug is of critical importance in instigating an immune response, which relates to the concept of “getting things just right” [13, 57]. For instance, administration of cyclophosphamide on a 6-day repeating schedule (Q6D) at 140 mg/kg per dose, dramatically improves the therapeutic outcome for murine GL261 gliomas through immunomodulatory mechanisms [9, 10, 11]. Other drug treatment schedules, however, did not result in the same efficacy. This loss of efficacy correlated with reversal of an initial anti-tumor immune response, despite ongoing ICD drug treatments [9]. Intriguingly, cyclophosphamide treatment of Lewis lung carcinoma (LLC) and B16F10 tumor at the same dose and on the same Q6D schedule did not result in tumor regression or immune cell recruitment, despite the intrinsic sensitivity of these two tumor lines to activated cyclophosphamide [12].

It is clear that much remains to understand about the underlying mechanisms of ICD action including the impact of chemotherapy dose and schedule on the many factors linked to the ICD response [13]. In particular, we explore the use of well-designed mathematical models, which can help elucidate the complex interplay between the various players, making these models an invaluable complementary tool to *in vivo* experimental results for designing better treatments.

### 1.3 Mathematical modeling of tumors and the immune system

There is a rich tradition in utilizing mathematical biology to study cancer chemotherapy and the immune system. A large variety of models have been proposed, notably in the work of de Pillis and collaborators, who introduced a series of models that depict many of the interactions between chemotherapy and immunotherapy drugs, the immune system, and tumor progression [58, 59]. Closely related to our topic, Ciccolini et al. [60] proposed a pharmacokinetics and pharmacodynamics (PKPD) model for metronomic chemotherapy using gemcitabine, one that considers the effects of cytotoxicity on endothelial cells and the emergence of drug resistance. Ledzewicz, Behrooz, and Schättler proposed a minimally parameterized mathematical model for low-dose metronomic chemotherapy that explicitly considers tumor vasculature [61], and in subsequent work [62] applied optimal control theory to this system, so as to devise a treatment schedule that can minimize tumor burden subject to appropriate constraints. To the best of our knowledge, only one study has looked at modeling the immune recruitment by ICD drugs [57]. In that work, however, there was no experimental validation of the model proposed.

Numerous cancer models have been proposed to account for the emergence of therapeutic resistance due to cancer cell heterogeneity. See, for example, the extensive references in [63]. In contrast, to our knowledge, no previous work has systematically and theoretically modeled what we call “systemic drug resistance,” by which we mean resistance as an immune-mediated dynamical phenomenon.

Here, we propose a mathematical model that is fit to experimentally observed tumor growth curves in GL261 tumor-bearing SCID mice that were given metronomic chemotherapy at Q6D to Q12D drug administration regimens [9]. The model proposed incorporates immune cell recruitment, as well as pharmacokinetics of cyclophosphamide. A fixed set of model parameters, not adjusted for individuals nor for drug schedule, is used to fit experimental data across these various drug regimens. To further validate the model, we not only compared the experimental fits to the measured tumor volume data, but also asked if the “latent variables” in our model, and which represent immune activation, match a second set of experimental data, not used in learning the model. Specifically, we asked if peak immune activation times in our model correspond to experimentally measured times. Finally, we investigate how our model can be used as a tool to identify better treatment schedules, build a quantitative understanding of the mutual interplay between drug, tumor, and immune system and to better predict drugs that induce immune cell recruitment in a clinical setting.

## 2 Materials and Methods

### 2.1 Background on cyclophosphamide and experimental results

#### 2.1.1 Summary of experimental data

The experimental data used in developing our mathematical model were derived from our previous work, where we implanted cultured GL261 gliomas cells in scid mice [9]. Tumors were allowed to grow to 300 to 1000 mm^3^, at which point the mice were treated with cyclophosphamide (CPA), given on different metronomic schedules. Greatest tumor burden reduction was observed when repeated doses of 140 mg CPA/kg-BW (body weight) were administered every 6 days. Comparisons were made between the every 6-day schedule and three other schedules: treatment every 9 days; treatment alternative between every 6 days and every 9 days; and treatment every 12 days. Additionally, we examined a dose of 210 mg CPA/kg-BW given every 9 days, which matches the overall dose of schedule of the every 6-day schedule. Tumor growth curves were reported for drug-free controls, and for the following regimens: a single CPA administration given at day 0 (1-CPA), 2 CPA treatments given at days 0 and 6 (2-CPA), as well as 3 CPA treatments given at days 0, 6, and 12 (3-CPA). Our previous work also reported relative gene expression levels for immune cell markers for NK cells, dendritic cells, and macrophages for the 1-CPA, 2-CPA and 3-CPA treatment regimens. There were *n* = 4 − 12 tumors per treatment group.

From our published data [9], the data indicates the host immune system takes 6 to 12 days before it starts to significantly impact the tumor growth. From that observation, we infer that short-term slowdown in tumor growth, within 1 or 2 days of an injection, is likely caused by cyclophosphamide-induced tumor cell death. We observed a decrease in immune cell number immediately after drug administration, highlighting the drug’s cytotoxic effect on immune cells as well as cancer cells. Notably, the chemoattractant CXCL10, which acts on most innate immune cells, and which is induced by IFN-*λ*, increased following first CPA injection, peaking around six days post administration. Between six and twelve days post injection, there is an increase in other innate immune cells markers, such as Nkp46, Nkg2d, Prf1, Gzmb for NK cells, Cd207 and Cd74 for dendritic cells, and Cd68 and Emr1 for macrophages [9].

Based on the impact of various treatment schedules tested on tumor volume, it is apparent that 12-18 days after the CPA treatment is halted, the tumor will cease to shrink and then starts to regrow. Based on treatments where the same dose of CPA is repeated at regular intervals, we conclude that a 6 day break between CPA doses at 140 mg/kg is ideal for maintaining prolonged immune cell recruitment as well as tumor shrinkage. Increasing the number drug-free days between treaments from 6 to 9 to 12 days increased the number of tumors that escapes from drug-induced tumor regression and also shortened the interval prior to tumor growth rebound [10]. Further analysis of tumor infiltrating immune cells in these models indicated there is a a strong correspondence between loss of immune response and tumor escape. Another study indicated that breaks in drug treatment shorter than 6 days also led to worse performance for the same AUC (area under the curve for the administration of the drug) with noted absence in immune recruitment if CPA dosing were given to the mice daily [10].

### 2.2 Modeling the immune-tumor-drug interactions

A schematic of our model, highlightling the main interactions between tumor, drug, and the immune system, is shown in Fig. 1.

**Figure 1:**
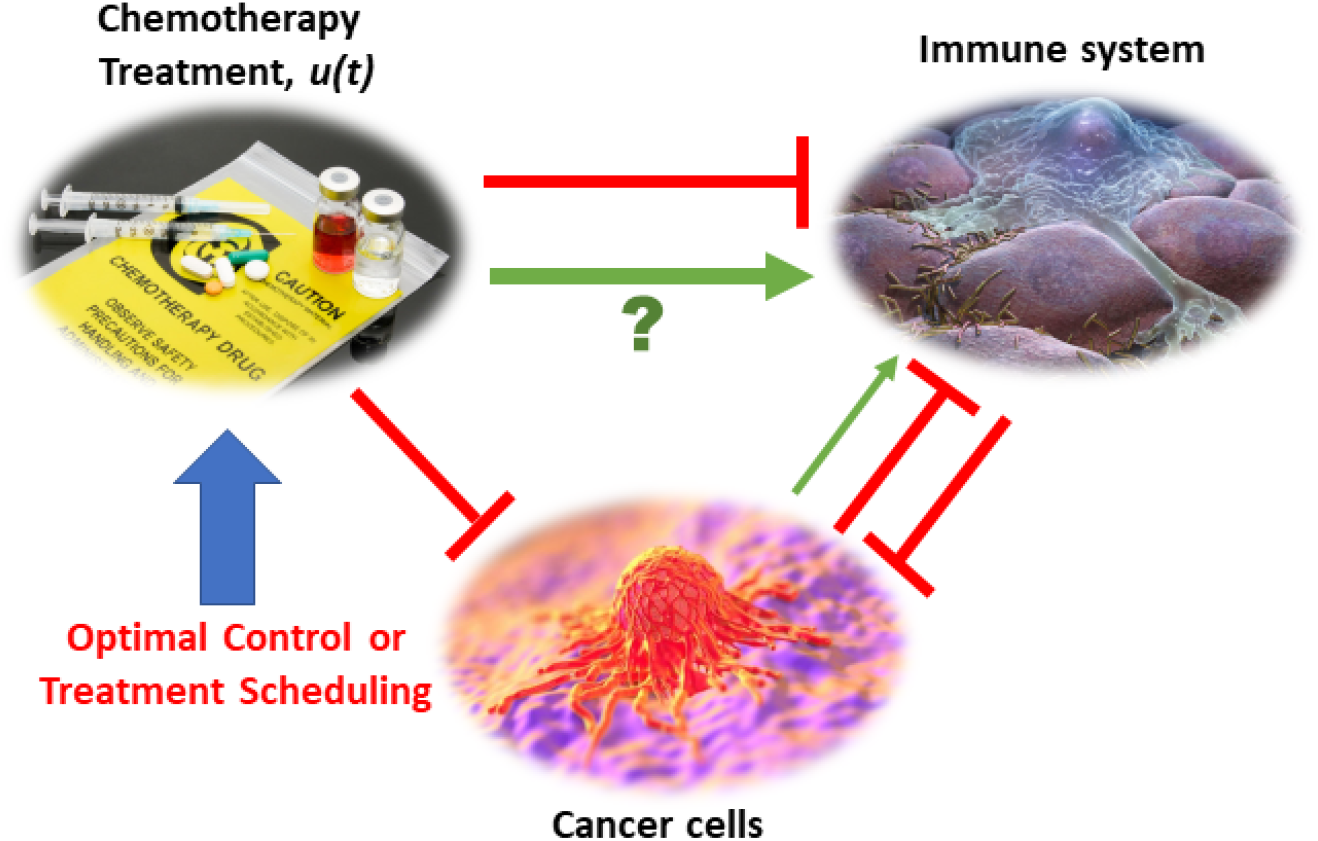
Mobilizing and sustaining a strong immune response require striking the right conditions of drug dosage and duration of brreak between drug administrations. The classical view of chemotherapy treatments is provided by the red arrows indicating inhibitory effects in which the chemotherapeutic drug suppresses both the immune system and the tumor, while the tumor and immune system repress one another. In this view, the immune system’s ability to act on cancer cells is greatly reduced by repeated cytotoxic drug administrations. The immune system can also be recruited to act on cancer cells by, for example, antigen presentation. The focal point of this work is to improve the mechanistic understanding on how applying the right metronomic chemotherapy regimen can, in specific cases, stimulate a large increase in the immune response (indicated by the green arrow with the question mark). Both the immunosuppressive and immunostimulatory effects of the drug are considered in our model, with the coexistence of these two seemingly opposite effects being an important point of emphasis.

Cyclophosphamide can be toxic to the immune system, as confirmed by experimental data showing that, one day after cyclophosphamide administration, there is a singificant decrease in expression marker genes for NK, DC, and macrophage cells in the tumor compartment [9]. When high doses of ICD drugs are used, toxicity to the immune system is so high that the role of the latter is greatly diminished. Given this finding, it is not surprising that traditional MTD chemotherapy not only has substantial side effects on the patient’s health, but also leads to immunosuppression and increases the risk of tumor relapse due to drug resistance. This then naturally leads to the question of how to represent (in a concise but not oversimplified mathematical way) the immunostimulatory and immunosuppressive effects of drug treatment when using a metronomic regimen of an ICD drug.

The paradoxical effect in which a drug reduces immune cell counts in the short term, while also enhancing the immune system in the longer term, is an instance of an “incoherent feed-forward loop” (IFFL). A similar paradoxical effect is that of the effect of treatment on cancer growth: on the one hand, the drug directly attacks the tumor, but on the other hand, through “friendly fire” also attenuates immune activity, thus degrading the anti-tumor response. IFFLs constitute one of the core network motifs in systems biology [64], and are found in processes as varied as gene regulation, immune recognition, synthetic biology, and bacterial motion [65, 66, 67, 68, 69, 70].

Some of the most common forms of IFFLs are illustrated by IFFL-I and IFFL-II block diagrams in Fig. 2. In our context, two forms of IFFL, promotion of an immunostimulatory intermediate (IFFL-I) or “repressing the repressor” (IFFL-II), are likely scenarios, and could occur in conjunction. Accordingly, to represent the immune system recruitment behavior, the mathematical model must include an intermediate that leads to immune cell recruitment in the longer term. The possible mechanisms are numerous, as summarized in the Introduction, and it is likely that the observed tumor growth is the result of both a repressor being repressed and some immunostimulatory element. Due to the lack of extensive immune data measurements, we only considered the immunostimulatory pathway in this work.

**Figure 2:**
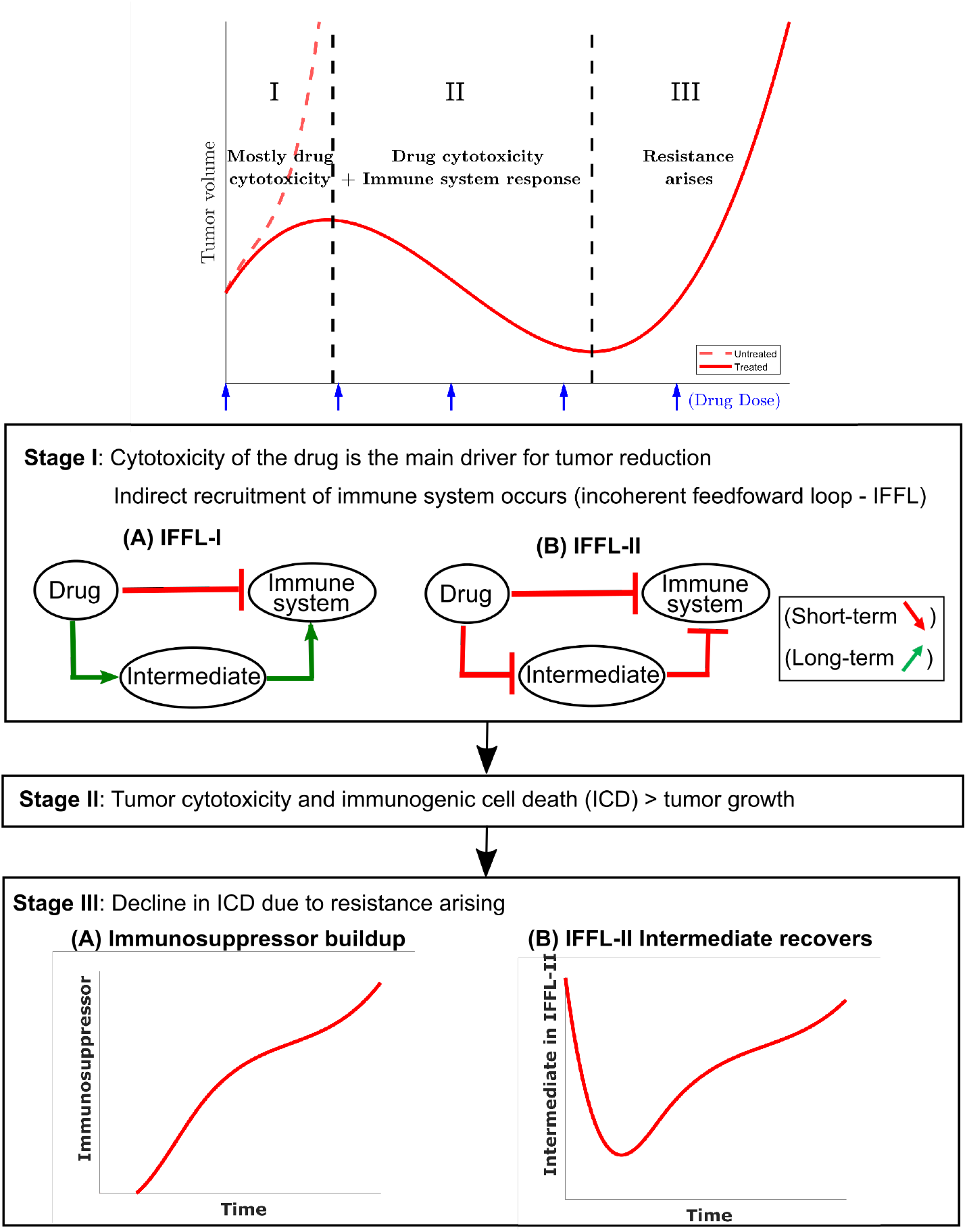
An illustrative tumor growth curve under metronomic chemotherapy treatment with repeated doses is shown and broken down into three representative stages. In stage I, tumor growth is primarily inhibited by the direct tumor cell cytotoxicity of the drug. There is typically a delay in the immune response due to the immune cell cytotoxicity of the injected drug. The immune cell data in [9] indicate that an immune response is weakened immediately after drug administration, but eventually recovers and leads to a strong response about 6 to 12 days later. Two possible incoherent feed-forward loop (IFFL) motifs that capture these short-term and long-term dynamics of the drug-immune system interactions are shown. In stage II, the tumor volume is typically reduced as the induced immune response becomes the principal contributor to tumor cell death. Stage II lasts until the induced immune response fades due to the emergence in drug resistance. Stage III typically has the tumor recovering its ability to grow in exponential fashion. Two possible scenarios are shown with the occurence of an immunosuppressor build up or the recovery of the intermediate in an IFFL-II scenario.

**Figure 3:**
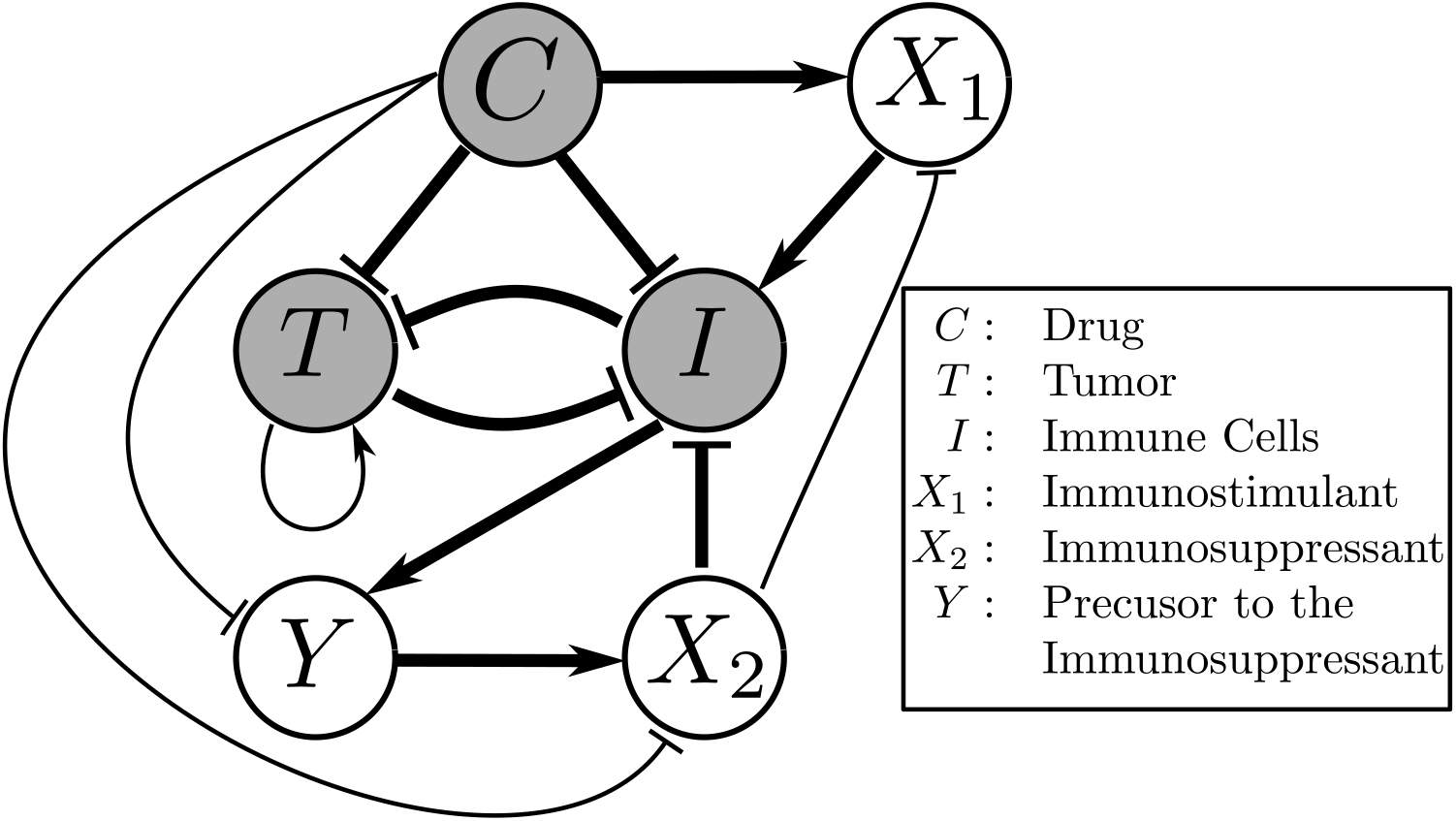
A schematic diagram of the interactions in our model. Thicker arrows represent well-known effects. Thinner arrows are hypothesized to be present, and also arise from our numerical fits to data.

In the event of tumor escape depicted conceptually in the bottom section of Fig. 2, the experimental data in [9] suggests that the immune system, which was reinforced through the first few injections of cyclophosphamide, fails to maintain the immune cell recruitment that allowed the host to keep the tumor at bay.

### 2.3 Mathematical model

The full mathematical model is given by a coupled system of seven ordinary differential equations (ODE’s). The first equation describe the change in CPA concentration over time (drug pharmacokinetics):

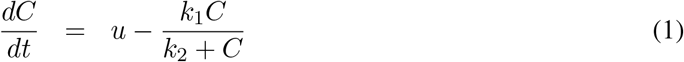

and the remaining five describe the effect of CPA on cancer and immune cells (pharmacodynamics):

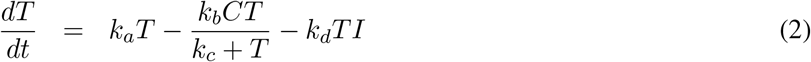

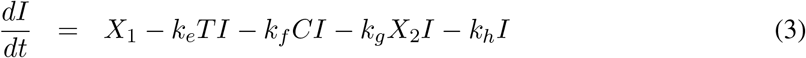

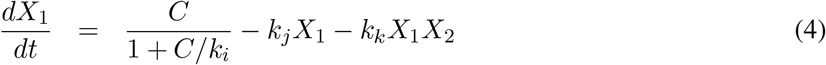

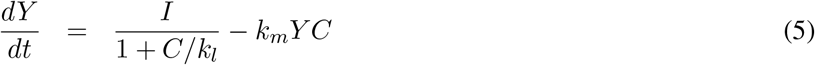

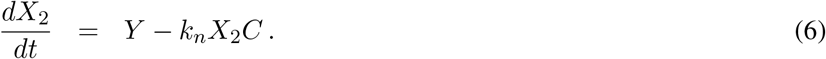

In the next section we describe in detail the various terms of the model, including the role of the variables *C* = *C*(*t*)*, . . ., X*_2_ = *X*_2_(*t*), which represent time-dependent changes of drug, tumor, and immune components, as well as provide descriptions and interpretation of the parameters *k*_1_*, . . . k_n_*. The input variable *u* = *u*(*t*) is introduced to account for the drug injection. The equation terms with missing parameters are the result to a nondimensionalization step that is detailed in the Appendix A.1.

#### The PK submodel

Equations (1)-(2) describe the change over time in concentration of cyclophosphamide after administration. This one-compartment model assumes that the drug is administered at a time-dependent rate *u*(*t*), resulting in a concentration *C*(*t*) in this compartment. This compartment is assumed to be where the interactions between the drug and its targets are assumed to take place. The drug is assumed to be cleared at a Michaelis-Menten (saturated) rate 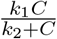.

The parameters *k*_1_ and *k*_2_ are obtained as part of the global fit to experimental data described later.

Our simple PK model is phenomenological, and intended to capture the delay in drug activity with respect to various cell types. Although the steps of CPA activation by liver cytochrome P450 metabolism have been well studied, including the pharmacokinetics of CPA in mice [71, 72], details of how the intermediates interact in real time with the immune system and the tumor cells remain largely unknown. In using a one-compartment model, the fast dynamics of the drug reaching the blood stream and the decomposition of CPA into its metabolites are assumed to occur nearly instantaneously (captured by *u*(*t*)) when studying this system on the time scale of days.

#### Tumor and immune dynamics

The full model describes interactions between five variables. These represent the tumor volume, denoted by *T* (*t*), the immune response denoted by *I*(*t*), and three phenomenological variables: an immunostimulatory intermediate species and an immunosuppressive intermediate species, whose counts are denoted by *X*_1_(*t*) and *X*_2_(*t*) respectively, and a precursor to the immunosupressive species whose count is denoted by *Y* (*t*). All these variables except *T* (*t*) are phenomenological representations of complex underlying phenomena, lumping together both cellular populations and chemical signals such as cytokines, and thus do not carry any tangible units. All three of these species are directly affected by the concentration *C*(*t*) of cytotoxic drug in the second drug compartment.

We assume that tumor volume grows exponentially, at a rate *k_a_*. In contrast to other mathematical models, we do not introduce a saturation term for tumor growth, because the data used in fitting is from mouse models, where a maximum tumor volume (or carrying capacity) is never reached for humane reasons. In our model, tumor cells can be killed by different two mechanisms: directly by the drug *C*(*t*) at a rate *k_b_*, and indirectly by the interaction between drug, tumor, and cytotoxic immune cells *I*(*t*) at a rate *k_d_*. The ratio 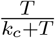 expresses that the cytotoxic death term is proportional in *T* when the tumor is small, but it eventually plateaus out when *T* becomes much larger than *k_c_*.

Taking into account these mechanisms results in the following equation for change in tumor growth rate:

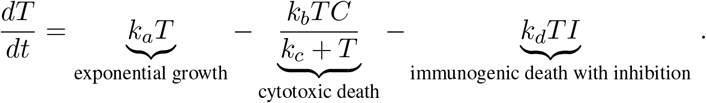

For the purposes of this analysis, we do not differentiate between various types of cytotoxic immune cells nor do we distinguish the effects of these cells from other immune factors such as chemokines or cytokines; instead, we track the change over time of an aggregate immune indicator *I*(*t*). We assume that this indicator increases through direct interaction with immunostimulatory intermediate *X*_1_(*t*) (which will be described next), and can either be inactivated through interaction with tumor cells at a rate *k_e_*, can decrease (“or die if seen as immune cells”) due to exposure to drug *C*(*t*) at a rate *k_f_*, can become suppressed through interaction with *X*_2_(*t*) at a rate *k_g_*, and can decrease at a natural rate *k_h_*. The term −*k_e_TI* represents both the activation *k_e_*_+_*TI* and inactivation of the immune response by tumor cells − *k*_*e*−_*TI* with the assumption that −*k_e_TI* = *k_e_*_+_*TI* − *k*_*e*−_*TI*. Through the result of fitting, we found that defining *k_e_* in this way led to a positive *k_e_* value.

This results in the following equation for change in the indicator *I* over time:

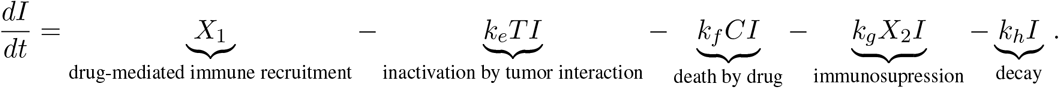

In addition to tracking the dynamics of tumor and immune cells, we introduce three phenomenological variables that are both affected by the drug, and can in turn affect both tumor and immune cells.

Firstly, we introduce an immunostimulatory intermediate *X*_1_(*t*), which impacts immune cells recruitment. We assume that it can be increased by an ICD type drug, such as cyclophosphamide, according to saturating function 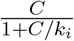, where immune cell recruitment is linear up to a threshold *k_k_*, and becomes saturated when *C*(*t*) *> k_i_*, representing an upper bound of drug-induced immune recruitment. The immunostimulatory intermediate *X*_1_(*t*) is assumed to decay at a rate *k_j_*, and to be inactivated through interactions with an immunosuppressive factor *X*_2_(*t*), which will be defined next. The resulting equation for change over time of immunostimulatory factor *X*_1_(*t*) is as follows:

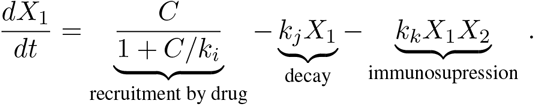

Finally, we introduce two immunosuppressive factors *Y* (*t*) and *X*_2_(*t*) that impact tumor-immune dynamics and their interactions with the drug *C*(*t*). They are described by the following two equations:

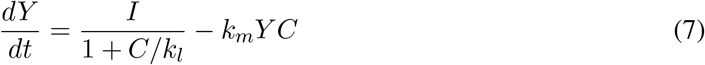

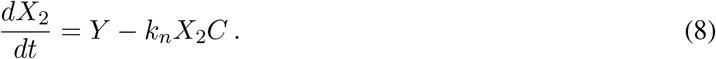

The two compartments are introduced to account for the significant delay between the effect of the drug on immunostimulatory vs immunosuppressive arms of the immune system. The immunosuppressive precursor *Y* (*t*) is assumed to be induced by *I* and a effectiveness term 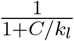 that is mediated by the drug *C*, and can be removed through interaction with the drug *C*(*t*) at a rate *k_m_*. The remaining precursor *Y* (*t*) increases formation of the immunosuppressive intermediate *X*_2_(*t*), which can also be damaged by the drug *C*(*t*) at a rate *k_n_*.

Equations (7)-(8) account for cytotoxic effects of cyclophosphamide not only on cytotoxic immune cells, such as NK and CD8+T cells, but also on immunosuppressive cells, such as MDSCs and Tregs [73, 74, 56]. The immunosuppressive intermediate *X*_2_ is vital to the appearance of systemic drug resistance when the treatments are continuously repeated over long treatment periods.

## 3 Results

In this section, we first use the phenomenological model described above to fit experimentally observed tumor growth curves. We then validate the model by comparing model predictions for the immune compartment to experimental data; notably, the data for the immune compartment were not fit, and thus provide an independent validation of the model, where a single set of parameter values was sufficient to recapitulate experimentally observed dynamics. Finally, the validated model is used to make predictions about treatment regimens that have not been tested experimentally.

Here, the notations 1-CPA, 2-CPA, 3-CPA are used to indicate that 1, 2 or 3 doses of CPA are given 6 days apart. The first dose is always given on day 0. CPA/6-days, CPA/9-days, CPA/12-days indicate that treatments were given 6, 9, or 12 days apart, respectively. CPA/9days(210mg/kg) indicates that the drug doses of 210 mg/kg (rather than 140 mg/kg in other treatment groups) were administered 9 days apart. Finally, the abbreviation CPA/6-9days indicates a break of 6 days between first and second doses, and a break of 9 days between second and third doses.

### Population fits to the experimental data

The experimental growth curves for individual tumors were obtained from [9] and fitted using the objective function outlined in Appendix A.2. The minimization of the error criterion was carried out using *fmincon* with the interior-point algorithm in MATLAB (Release R2019a, Mathworks, MA). The values of *a*, *b*, and *c* in the error criterion were chosen as 19.31, 1, and 227.6, respectively. These values were found to provide a balance between providing enough weights to errors involving small experimental values, while also ensuring that the desired phenomena of tumor evasion and immune recruitment are captured appropriately by the model.

The initial values of the state variables were assumed to be 0 except for tumor volume. The rationale behind this assumption is detailed in Appendix A.2. The initial guesses to the optimization problem used for finding population parameter fits were drawn from a uniform distribution on a log-scale to sample several orders of magnitude. Numerous initial guesses (> 10^5^) were tested using the Northeastern Discovery computer cluster. Due to the nonlinearity of the model, multiple sets of parameters yielded equally good minimal objective function evaluations. Of the 10 best evaluation values, the set of population parameters that best captured the tumor escape effects across various treatment conditions was picked and presented in Table. 1. Outliers were excluded using the procedure outlined in Appendix A.2.

**Table 1:**
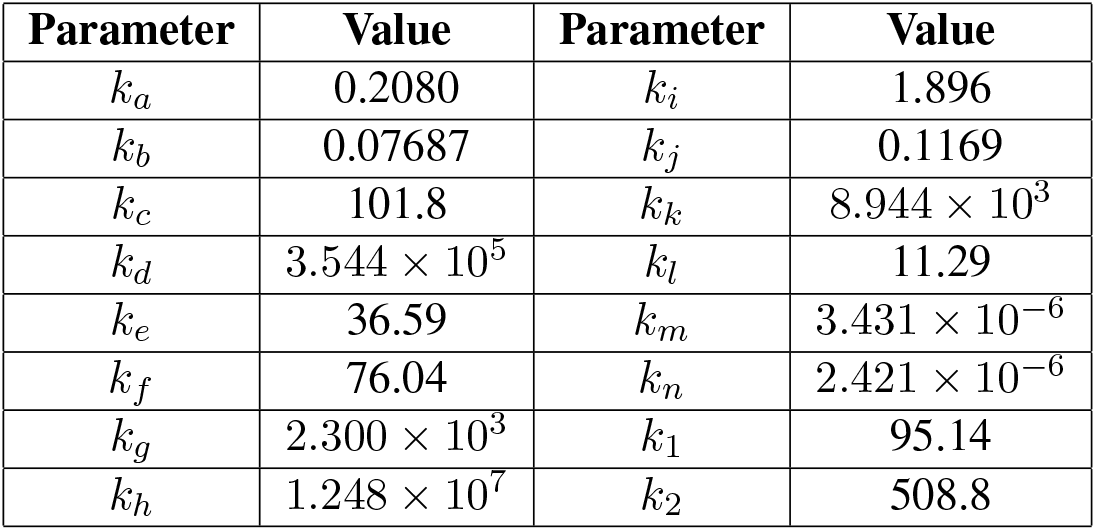
Fitted parameter values.

| Parameter | Value | Parameter | Value |
| --- | --- | --- | --- |
| $k_a$ | 0.2080 | $k_i$ | 1.896 |
| $k_b$ | 0.07687 | $k_j$ | 0.1169 |
| $k_c$ | 101.8 | $k_k$ | $8.944 \times 10^3$ |
| $k_d$ | $3.544 \times 10^5$ | $k_l$ | 11.29 |
| $k_e$ | 36.59 | $k_m$ | $3.431 \times 10^{-6}$ |
| $k_f$ | 76.04 | $k_n$ | $2.421 \times 10^{-6}$ |
| $k_g$ | $2.300 \times 10^3$ | $k_1$ | 95.14 |
| $k_h$ | $1.248 \times 10^7$ | $k_2$ | 508.8 |

In Figs. 4 and 5, the simulated growth curves generated using parameters summarized in Table 1 are shown side by side with the corresponding experimental data.

**Figure 4:**
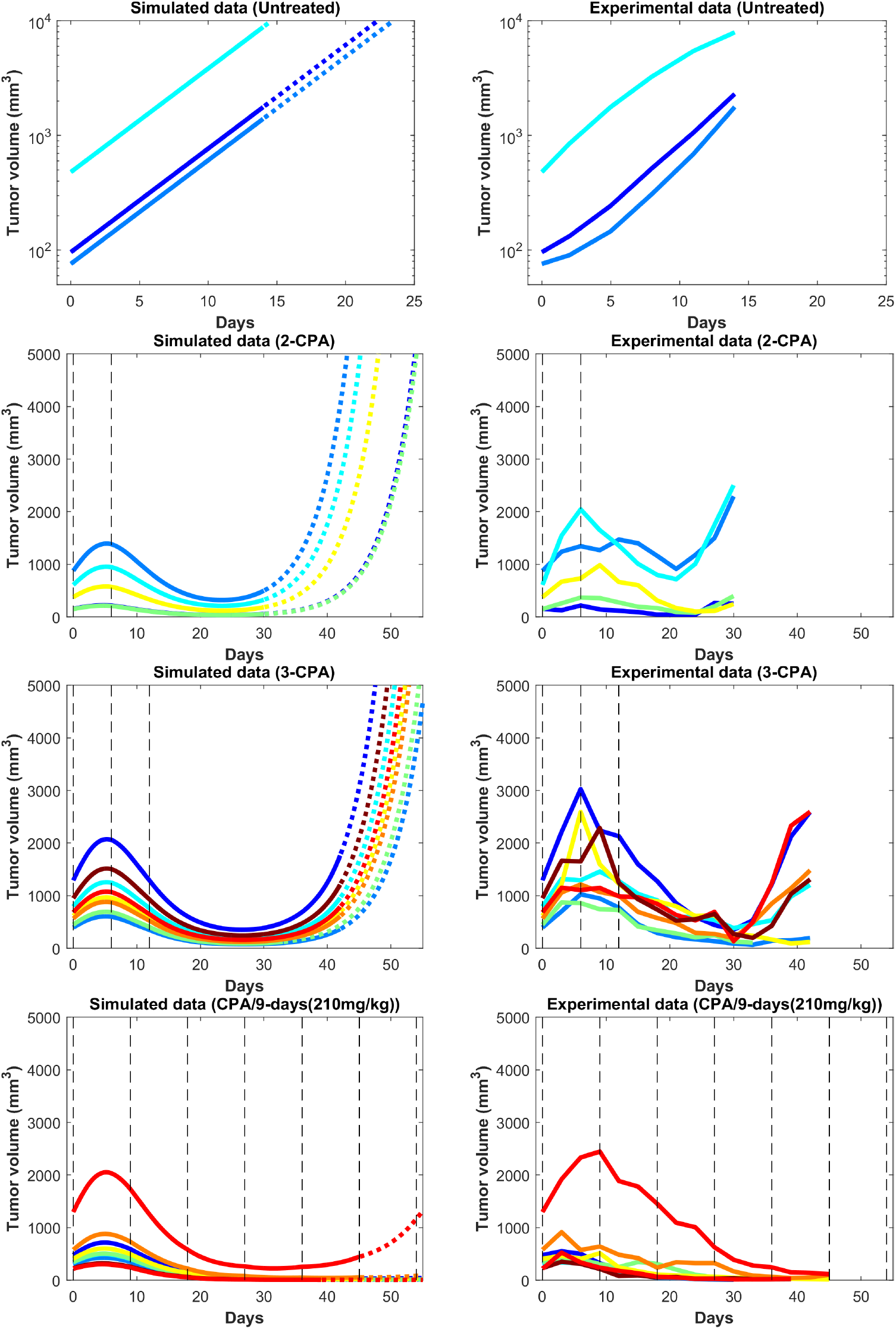
Simulated and experimental growth curves for the scenarios of 1-CPA, 2-CPA, 3-CPA and doses of 210 mg/kg given every 9 days (CPA/9-days/210). When not specified, CPA doses are 140 mg/kg. Black dashed lines represent the time at which drug treatments are given. Solid lines represent fitted data; dotted lines represent predicted growth curves extrapolated from the model.

**Figure 5:**
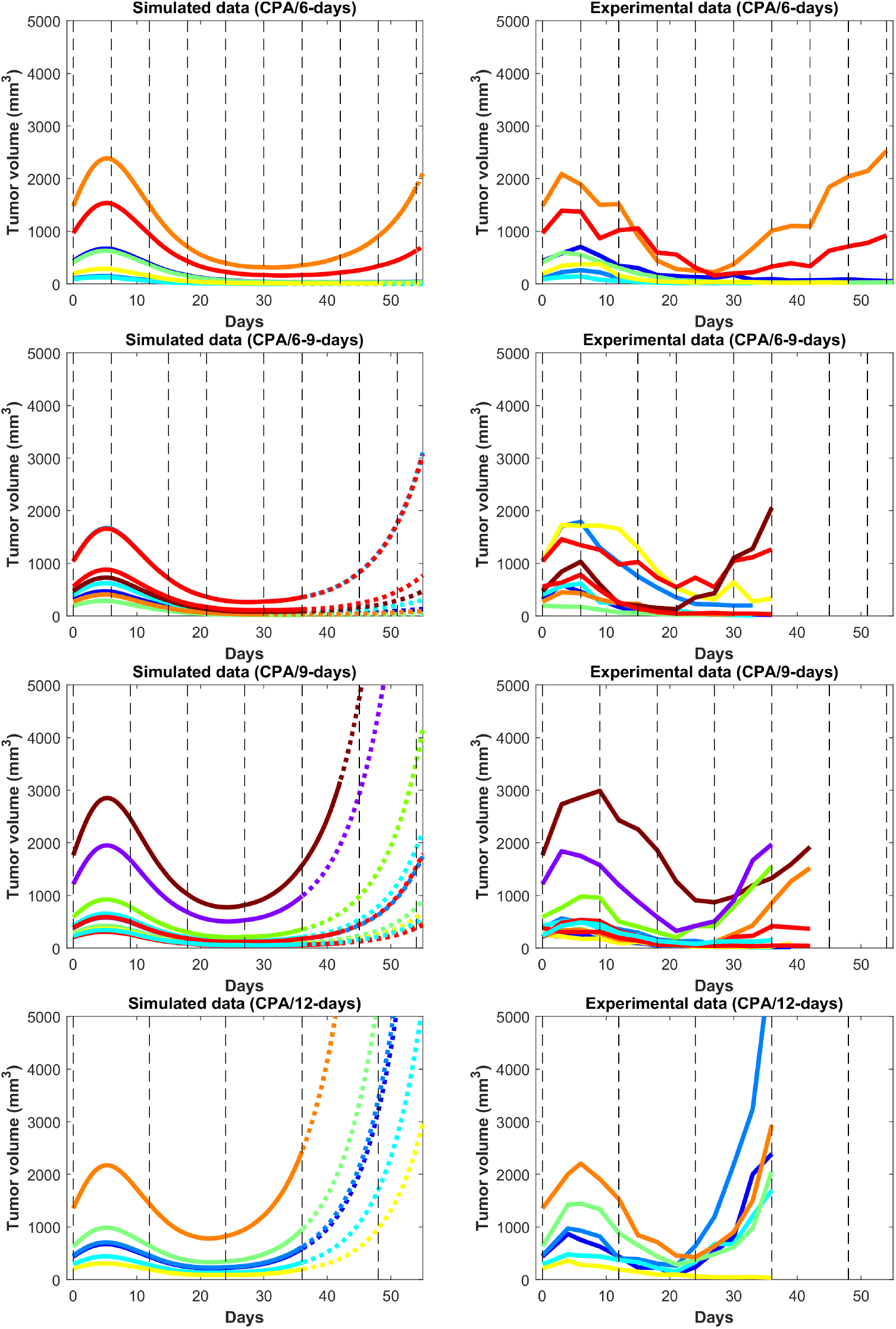
Simulated and experimental growth curves for the scenarios CPA/6-days, CPA/6-9-days, CPA/9-days, and CPA/12-days. Solid lines represent fitted data; dotted lines represent predicted growth curves extrapolated from the model.

Despite the use of pooled mouse data for curve fitting, there is a high degree of agreement between experiments and model fits, particularly for tumors that remained largely suppressed throughout treatment. The nature of rebounding (or escaping) tumors makes the observations very stochastic in nature, while the models are progressively transitioning between regressing and rebounding tumors under conditions as the initial tumor volume gets larger (an example of this transition being the CPA/6-9-days treatment). Allowing for small variations in the parameters between individual experiments is likely to account for such variability. The primary intent is to showcase that one unique set of population parameters can capture the qualitative and quantitative behaviors of a large set of different metronomic chemotherapy treatments that involves induction of anti-tumor immune responses and drug resistance.

The agreement between simulated and fitted experimental growth curves is high for the untreated, 2-CPA, 3-CPA and CPA-9days(210mg/kg) cases, as can be seen in Fig. 4. There is a discrepancy in between the model prediction and the experimental data for the largest tumor in the 2-CPA scenario. In the repeated treatments, the model captures well the progressive apparition of rebounding tumors as intervals between drug administration increase from 6 to 12 days in 4 increments. For the CPA/12-days scenario, the model predicts that tumor escape will occur up to ten days after when it actually occurred experimentally; nevertheless, the model is able to capture the qualitative effect of this treatment regimen, which fails rapidly within the first 3 drug injections.

The 1-CPA set of data was small in size (*n* = 5, before an outlier was excluded) and the tumors seemed to have been implanted with tumors growing at a significantly faster rate than the rest of the dataset. Given that each metronomic scenario was from a different batch of mice, systemic experimental variations in the fitted data can account for some of these observed discrepancies and can be hard to distinguish from model deficiencies. From Fig. 6, the three treatment conditions of 1-CPA, CPA/9-days and CPA/12-days are not expected to differ until day 9, but the plots indicate that the experimental data is inconsistent. In addition, 1-CPA and CPA/12-days have mice undergoing the same treatment conditions until day 12.

**Figure 6:**
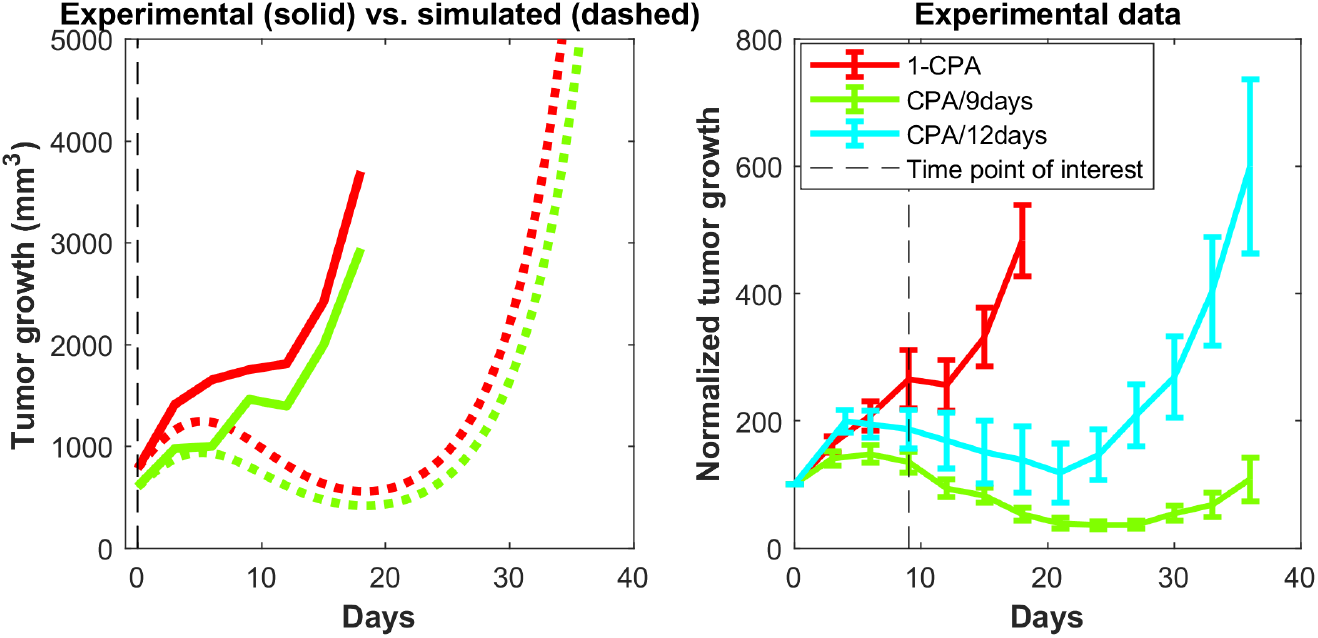
Discrepancies in modeling 1-CPA. On the left, are shown the experimental data and the simulated data, with the latter data predicting much slower tumor growth than was seen in the experimental data. On the right, the 1-CPA data is plotted using the normalized tumor growth and compared with CPA/9-days and CPA/12-days treatments. Note that the 1-CPA treatment is equivalent to the CPA/9-days and CPA/12-days data up to treatment day 9 and treatment day 12, respectively, barring systemic errors in the experiments.

In Fig. 7, the mean normalized tumor volumes are given for each of the conditions modeled and shown with the corresponding experimental data. There is a slight delay in the 2-CPA case when it comes to the rebounding behavior of the tumors. The CPA/12-days discrepancy in the time of rebound can also been seen. Notably, the deficiencies in the fits are more pronounced on the marginal cases of longer breaks and shorter treatments. It is possible that these edge cases require special attention of other phenomena that are discussed in [13], but that likely will require additional experimental data to model appropriately. Also shown in Fig. 7 is the difference between the normalized average fits of the fitted data without outliers and the experimental data with outliers. The trends were not affected by removing outliers with the exception of CPA/12-days that yielded large discrepancies between 4 and 25 days. This was due to two small tumors that were excluded as they grew very rapidly to be much larger than other tumors that had higher initial volumes.

**Figure 7:**
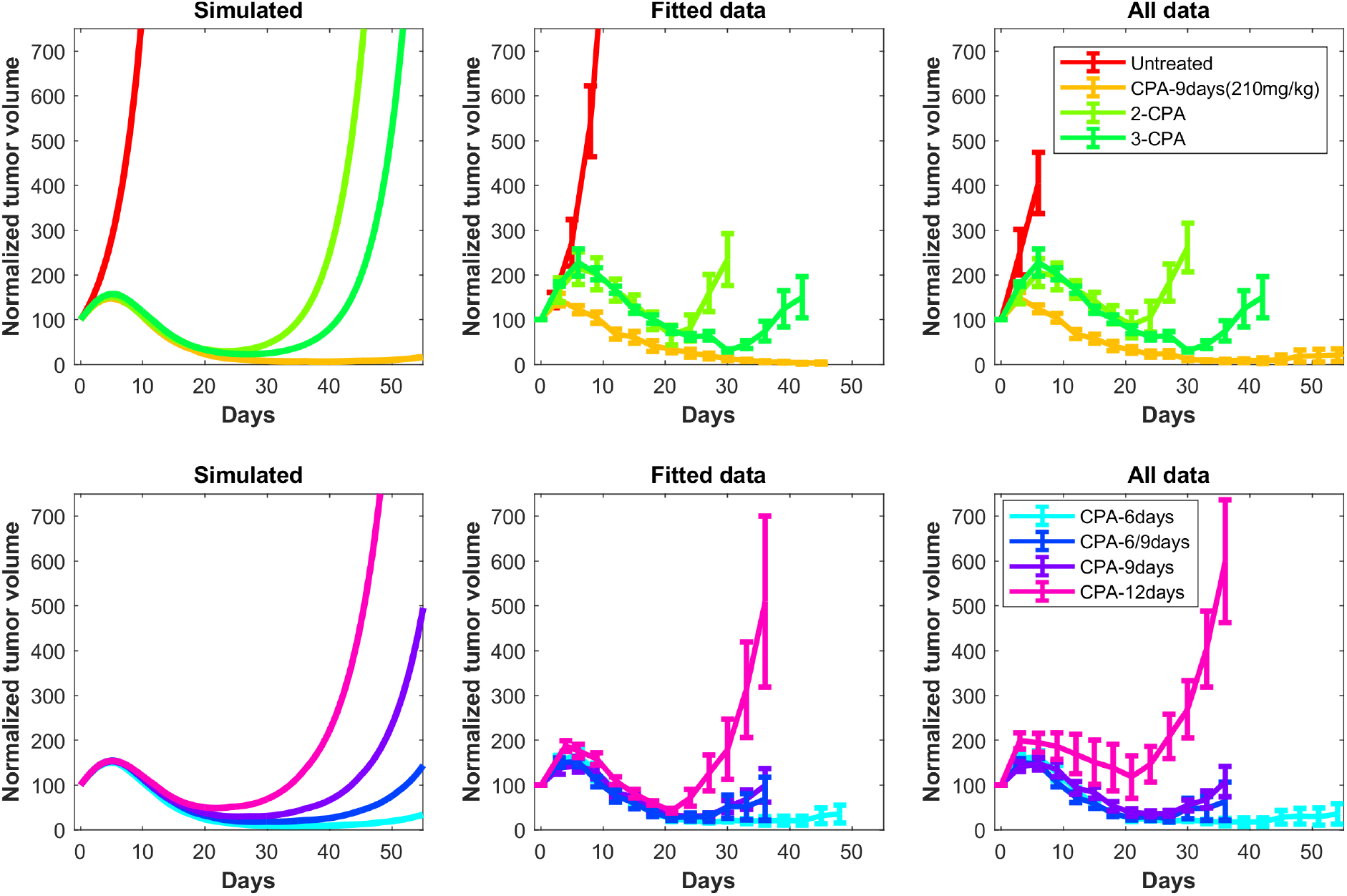
Simulated tumor growth curves of all the considered treatments scenarios were normalized such that the first time point of each curve has an initial value of 100. The mean of the normalized curves for each treatment condition was then calculated and plotted on the left panels. The right panels are the original experimental data from [9]. These data also show the difference between the normalized average fits of the fitted data without outliers (middle panels) and the experimental data including outliers (right panels). The outliers are highlighted in Appendix A.2

### Predictions regarding the immune system

In Fig. 8, predictions are made for 1-CPA, 2-CPA, 3-CPA for the immune system behavior. Notably, the immune data was never used in the model fitting, but the model predictions of the immune system corresponds well with that of the immune data in [9]. In the model, the immune system is assumed to be an aggregate of multiple immune cells and related factors. The experimental immune data, shown in Fig. 8, separates the data in two categories: the immune cell markers and the chemokine, cytokine, and adhesion molecule markers. The experimental data is very clear on indicating that new injections of the drug has a short term negative effect on the immune cell populations. However, this is not often the case for cytokines or chemokines, which can be recruited and remain at high doses, even after a second injection such as seen in the 2-CPA data. Considering both effects of the immune cell markers and cytokines and related molecules, the predictions of the immune system behavior by the model seem to be an aggregate of these responses. It estimates well the estimated immune cell peaks of 12 days, 18 days, and 24 days, respectively for 1-CPA, 2-CPA, and 3-CPA. There is a sparsity to note in the experimental data as only 4 time points were analyzed for each treatment condition.

**Figure 8:**
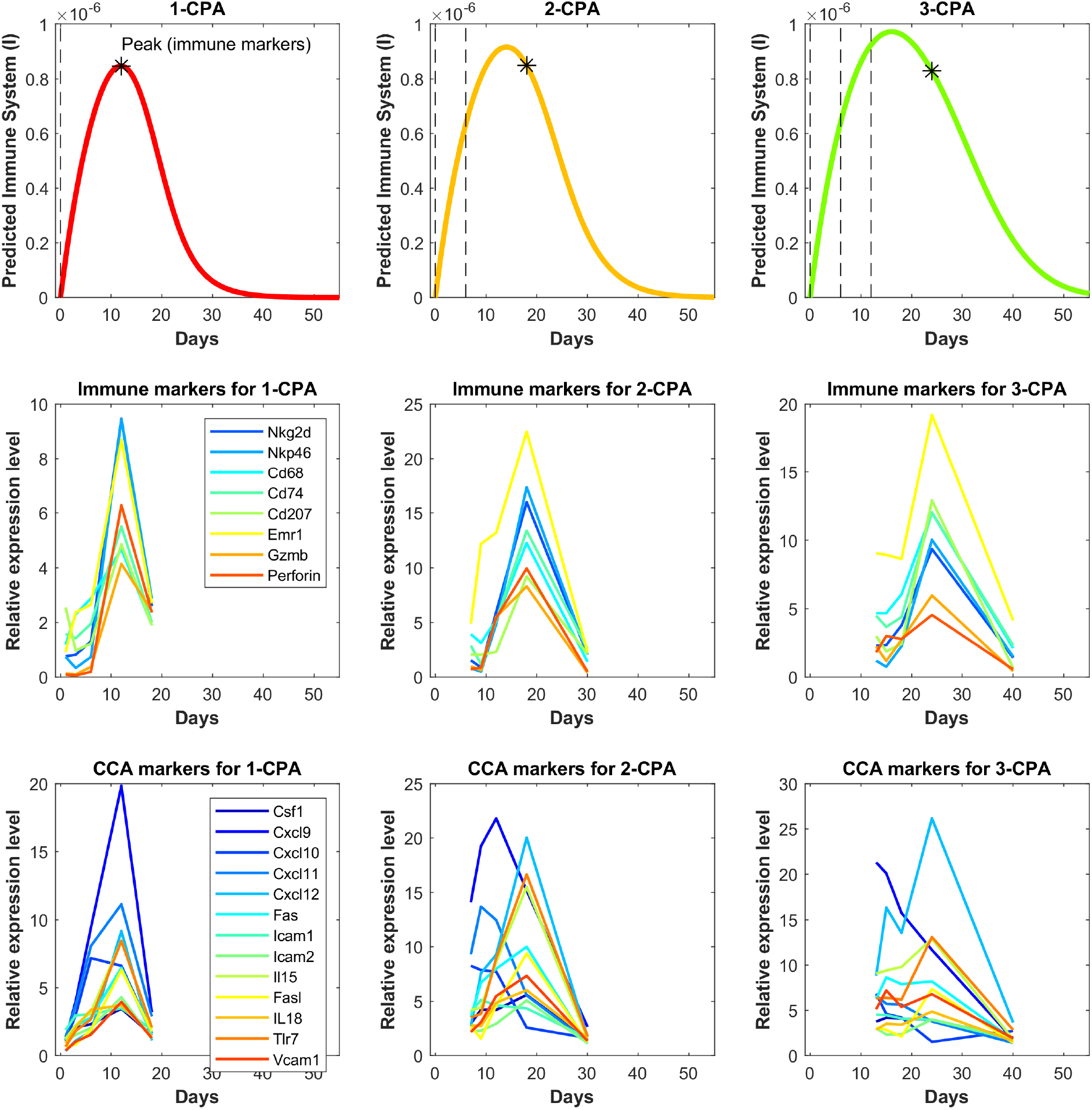
Independent model validation through comparison of model predictions to available immune cell data. The top row shows the predicted immune system recruitment by the fitted model for the 1-CPA, 2-CPA, and 3-CPA treatment conditions. The vertical black dashed lines indicate times of drug injections at 140 mg/kg. The initial volume for the simulated tumors yielding these curves was assumed to be 1000 mm^3^ at the time of the first injection. The middle row shows the experimental data of the gene expressions for various immune markers linked to the innate immune cells (macrophages, dendritic, and NK cells). Similarly, the bottom row shows gene expression data for various chemokines, cytokines, and adhesion molecules (abbreviated CCA on the plots). The data is ordered such that each column represents the same treatment condition.

### Predictions for CPA/9-6-days and CPA/7.5-days

The validated model was used to make predictions about the effect on tumor growth of dose administration regimens that were not experimentally tested. One such example is shown in Fig. 9, where drug is administered at alternating breaks of 9 days and 6 days (CPA/9-6days). Interestingly, the model predicts that CPA/9-6days is considerably inferior to the CPA/6-9days regime, suggesting that shorter breaks between drug administrations early on improve outcomes, as compared to longer breaks. CPA/7.5-days is a little better than CPA/9-6days, especially with smaller initial tumor volumes.

**Figure 9:**
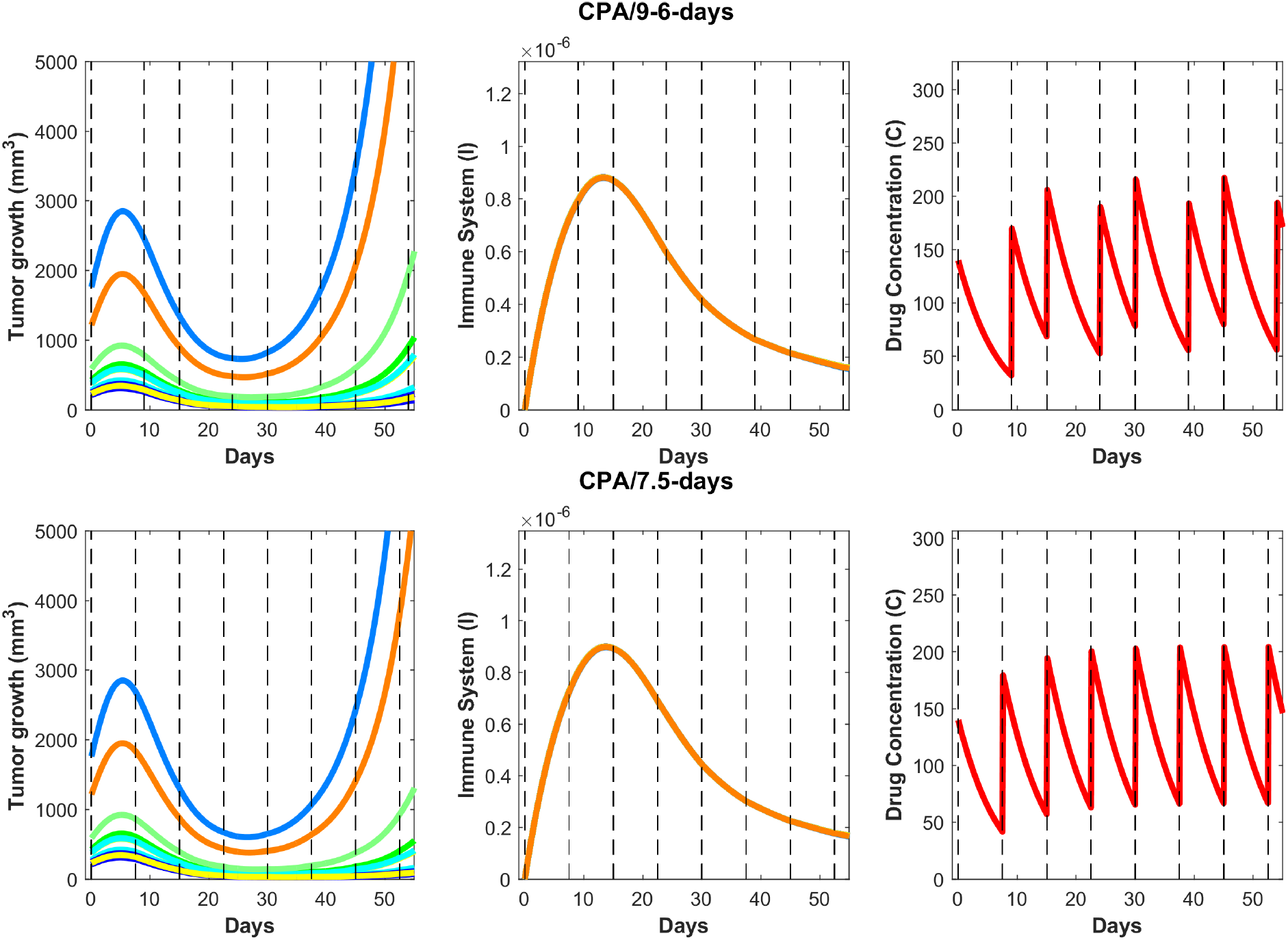
On the top row, predictions for the CPA/9-6-days drug regimen where the first break is 9 days followed by 6 days, then 9, then 6 and so on, alternatively. On the bottom row, similar predictions are made for CPA/7.5-days.

**Figure 10:**
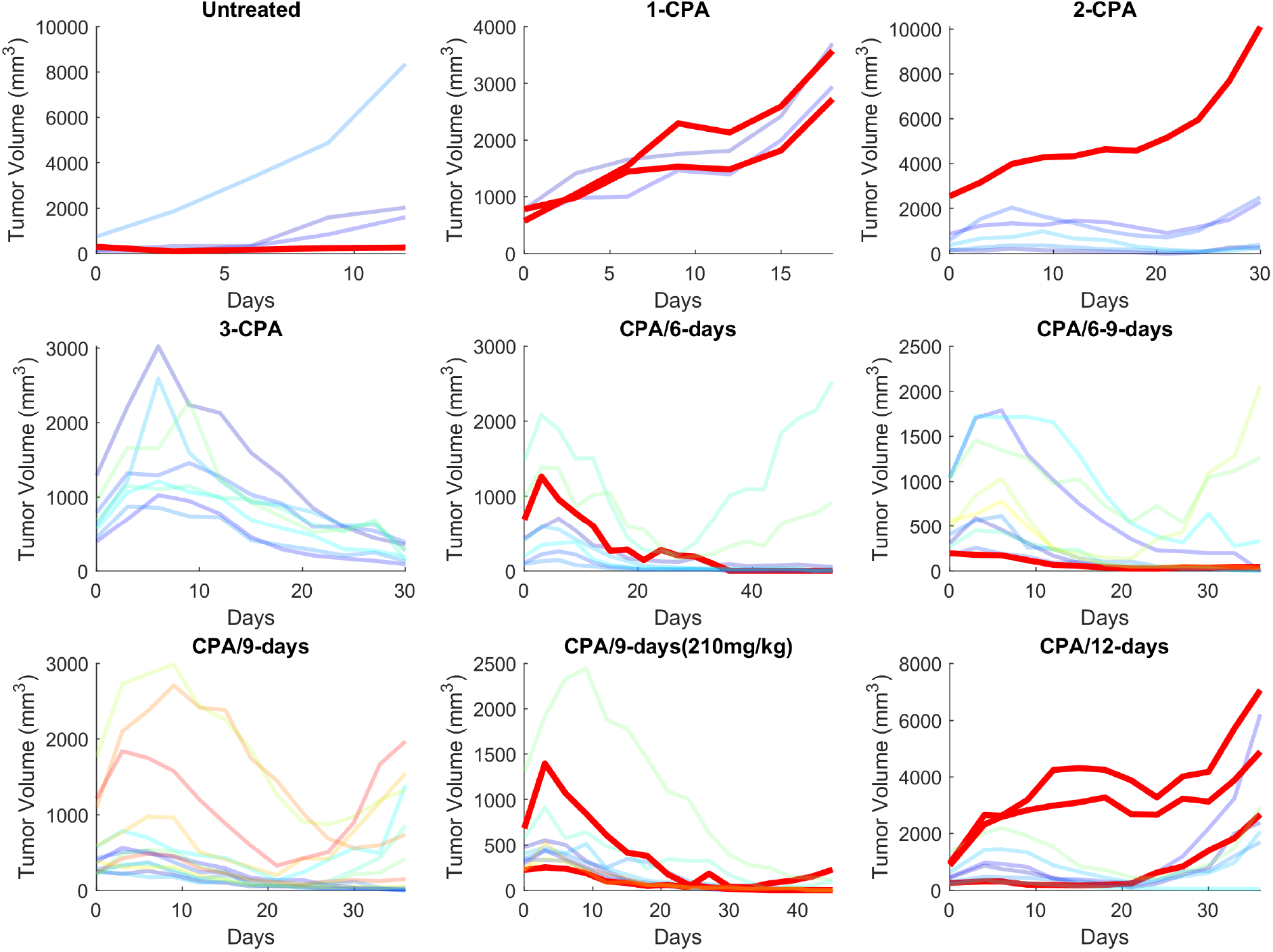
The experimental data in [9] are plotted with the outliers highlighted in red. Out of the 65 time series data considering 9 different treatment conditions, 11 were excluded in the fitting of the population parameters.

We hypothesize that shorter breaks early on allow sufficient tumor burden reduction to enable cytotoxic immunity to have greater impact on the smaller tumor. Notably, within this framework, the standard approach of maximizing tumor burden reduction would cause excessive damage to the immune system, so chemotherapy-induced tumor burden reduction should be sufficient to augment the effect of the immune system but not act to its detriment. Based on our analysis, although CPA/9-6days schedule (Fig. 9) should be superior to the CPA/9days schedule (Fig. 5), the two simulated datasets are almost identical in behavior. Upon closer examination of the cancer cell death due to activity of the immune system, as opposed to the drug cytotoxicity, the model indicates that even after the anti-tumor immune recruitment decreases, small tumors remain under control due to drug cytotoxicity. However, for large tumors, even a slight decrease in anti-tumor immunity can determine the difference between tumor regression and tumor progression. Looking at the immune response in Fig. 9, a slight decrease around 15 to 20 days after the first treatment leads to strong rebounding behaviors in the two largest simulated tumors.

## 4 Discussion

The administration of cyclophosphamide under MTD regimens can undermine the immune system’s ability to help control tumor growth. Changing the dose and timing of drug administration has been shown to restore the ability of cytotoxic immune cells to target tumor growth in glioma mouse models [9, 10], suggesting the existence of a “sweet spot” that can minimize damage to the immune system and maximize anti-tumor immune effects. To formalize the mechanisms that may underlie experimentally observed variation in response with respect to drug dose and schedule, we propose a phenomenological mathematical model that captures key processes that may underlie interactions between drug, immune system, and tumor. Through these efforts, we aim to identify key mechanisms that may give rise to a strong and sustained immune response, within the broader context of improving the understanding of cancer treatments that target not only cancer cells themselves but also the tumor microenvironment, and in particular, the immune system.

The proposed phenomenological model was fit using the method outlined in Appendix A.2 to generate a single set of parameters that was able to capture well tumor growth dynamics across nearly all experimentally tested drug treatments (Figs. 4 and 5). The model was further validated through predicting the dynamics of the immune cells that were not used in the fitting process. We found a strong correspondence between immune data and predictions, despite the simplified nature of the underlying model (Fig. 8).

We then used the validated model to predict the impact on tumor growth of an alternative schedule of CPA/9-6days as well as CPA/7.5-days that have not been tested experimentally (Fig. 9). The model predicts that shorter breaks between dose administrations early on lead to greater tumor burden reduction and improved anti-tumor immunity; however, as was shown in [9, 10], the breaks cannot be too short, which in turn may lead to excessive immune cell depletion, not allowing cytotoxic lymphocytes time for replenishment. Therefore, the goal of “sweet spot” therapy is to reduce tumor burden sufficiently to allow cytotoxic immunity to persist and control the tumor.

Despite the undeniable complexity of the immune system, the proposed conceptual model of five differential equations coupled with a 2-dimensional PK model allowed us to capture tumor responses to various treatment schedules. While the model was used to understand and reproduce data that shows impact of variation in dosing and scheduling on tumor growth for a particular mouse tumor model, it also highlights the fact that that beyond simply understanding the interactions between the different components at play, a quantitative modeling approach may be able to help design better metronomic chemotherapy treatments. Furthermore, the conceptual nature of the proposed model enables us to pinpoint more specific mechanisms that are responsible for observed variations, which may not be possible with more detailed descriptive models.

Besides fitting well the experimental data on cyclophosphamide, the mathematical model developed here can be a valuable complementary tool to understand how drugs function and how they can be combined with other types of treatments, such as immunotherapies. It is also possible that drugs were discarded in the past given a lack of cytotoxicity may have immunogenic properties that could act as effective complements other treatments, a response that may have gone unnoticed due to MTD administration. Furthermore, the proposed framework suggests that what may appear as therapeutic resistance to the drug may in fact represent desensitization, a phenomenon that can be mitigated through alterations of the drug dosage and scheduling and through better understanding how these different components of the tumor microenvironment interact with one another, particularly immune cells and cytokines. This may open avenues to re-purposing existing drugs by appropriately altering the dosage and schedule of administration, an undertaking where quantitative approaches such as the one proposed here may prove to be indispensable.

We anticipate the proposed mathematical model will be useful for discovery of hidden potential of current drug treatments by building better quantitative understanding of the phenomena at play, and for designing more effective drug regimens that may not be intuitively apparent through exploring immunogenic potential of ICD drugs. The proposed model can also be used to investigate a variety of drug schedules and dosing regimens, as well serve as a building block for investigation of combination chemotherapy and immunotherapy treatments that will likely pave the future of cancer therapy.

## Acknowledgements

The work of Anh Phong Tran, M. Ali Al-Radhawi, and Eduardo D. Sontag was supported by NSF grants #1716623 and #1849588, while the work of David J. Waxman was supported by the NIH grant CA49248.

## Author Contributions

Conception and design: IK and EDS.

Development of methodology: IK, APT, MAA, and EDS.

Acquisition of data: JW and DJW.

Analysis and interpretation of data: All authors.

Writing, review and/or revision of manuscript: All authors.

Administrative, technical, or material support: EDS.

Study supervision: MAA and EDS.

## A Appendix

## A.1 Nondimensionalization of the model equations

Consider the equations of the model:

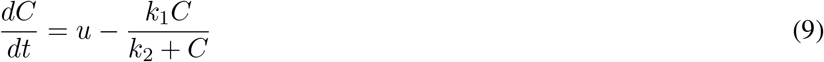

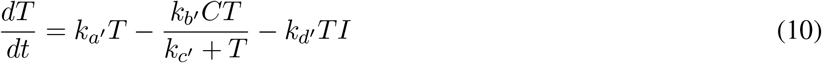

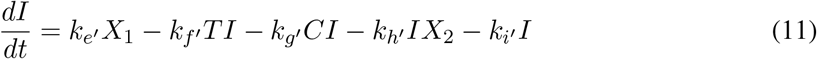

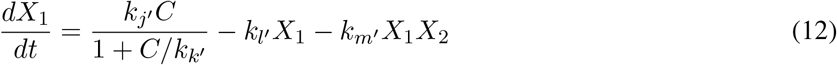

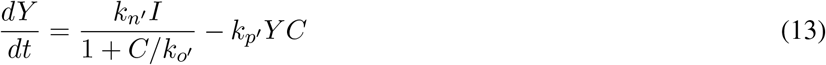

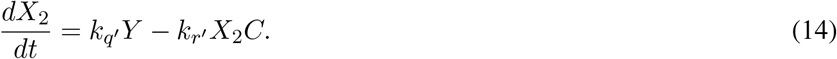

Substituting *I*, *X*_1_, *Y*, and *X*_2_ using the relationships:

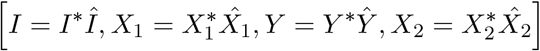

and limiting the scope to the equations for these state variables to be nondimensionalized yield the following set of equations:

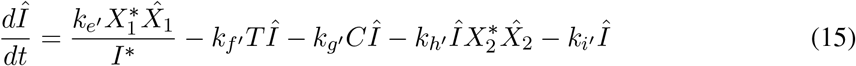

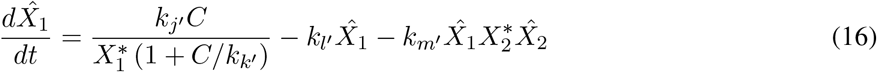

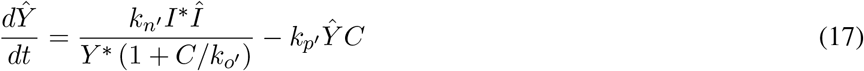

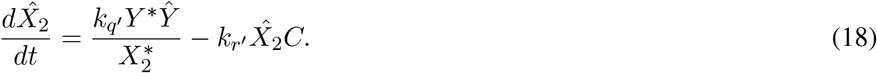

Making the following replacements

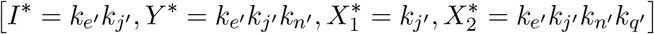

and rewriting the parameter names yield the following nondimensionalized set of equations with 4 less parameters:

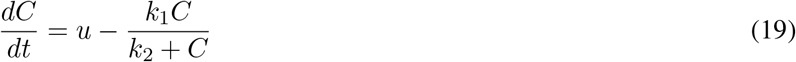

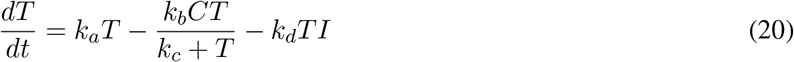

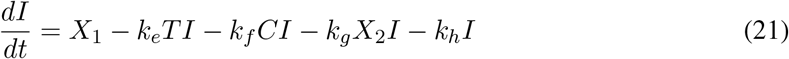

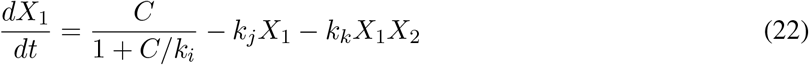

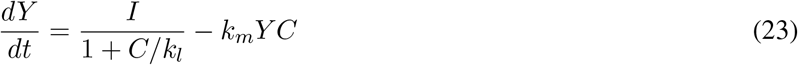

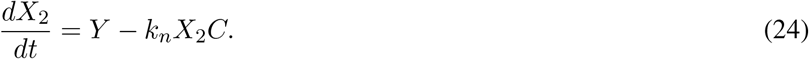

## A.2 Fitting methodology

## Error criterion

In this work, an objective function was defined for the nonlinear optimization problem that is used to fit the model parameters:

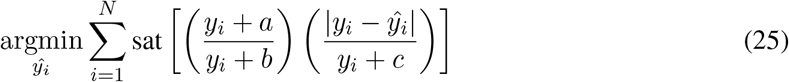

with *y_i_* being the experimental value and *ŷ*_*i*_ the predicted value. sat() is a saturation function values to 1. The values of *a*, *b*, and *c* are determined in such a way that low values of *y_i_* are not weighted too strongly. In the objective function, *b* and *c* can be interchangeable, so we will assume that *b < c*. In the limit that *b* < *y_i_* ≪ *a*, *c*, we get that:

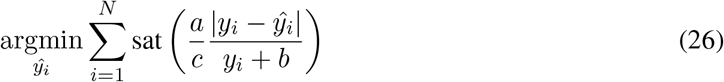

*a* and *c* are chosen such that 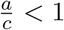 and b is small value that plays both the role of a regularizing term and avoids a division by 0. In the limit when *y_i_* is large, we get:

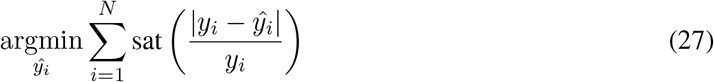

which is a standard normalization. The magnitudes of *a* and *c* play an important role in how quickly this limit of large *y_i_* is approached. Let’s note that if *y_i_* ≪ *b* ≪ *a*, *c* then the ratio becomes 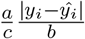. The parameters are chosen such that *bc* ≫ *a*, small values are filtered out, as these may be more susceptible to measurement noise or below a threshold of detection.

## Determination of the population fits

The main underlying assumption of the population fits is that all mice are characterized by the same parameters. In reality, there can be a myriad of ways mice can differ from one another. Notably, their immune system might be of different strength when fighting the tumor and different tumors can grow at different speeds. However, the immune data that was available [9] is confined to the 1-CPA, 2-CPA, and 3-CPA scenarios and only three individual mice per experiment. The tumor growth curves were thus the only data that was used to fit the model.

In addition to having the same parameters, the initial values of all state variables were assumed to be 0, except for the initial tumor volume. While the mice do have a still functioning immune system at the moment they are being monitored, an escaping tumor is a sign that the immune system is compromised or unable to contain the tumor growth. Thus, the underlying assumption is that the immune system at the start of treatment is either of negligible effect or its effect on the tumor growth is lumped inside the *k_a_* constant. Furthermore, the immune data from [9], also shown in Fig. 8, shows that the effect of cyclophosphamide given on a metronomic regimen can yield an order of magnitude increase in the gene expression of innate cell immune markers, and similar observations can be seen for the gene expression of other markers for cytokines, chemokines, and adhesion molecules.

The last assumption for the population parameters is that the input *u* appearing in Eq. 1 consists of a step input of 140 mg/kg in concentration, applied at the moment a dose injection. From the time-scale of the experiments, the time-scale of the drug injection is negligible when monitoring the tumor size every few days. The input and the state variables besides the tumor (*T*) are assumed to be unitless. For the drug regimen of CPA/9-days(210mg/kg), the step input for *u* was increased to 210 mg/kg in concentration for each dose injection in order to account for the higher dose.

## Handling the outliers

In this data, there are two types of outliers that were taken out of the analysis:

- Time series with average tumor volume above a given cutoff value of 2500 mm^3^ and that did not belong to the untreated group.
- Time series that contradicted the behavior of neighboring curves.

The first criterion led to the exclusion of 3 outliers out of 65 time series. Large tumors followed dynamics different from what the model could explain. However, the large tumor data is very sparse, so there did not seem to be enough data to confidently elucidate the functional form that governs these data points.

Given the interest in finding a set of population parameters that captured the quantitative and qualitative behaviors of the experimental data, a second criterion was used to avoid fitting the model with contradictory behaviors. To quantitatively measure these deviations, the experimental values of the tumor volumes at each time point at a given treatment condition are ranked. Defining *r_ijk_* that represents the rank for the time point *i* of time series experiment *j* at the treatment condition *k*. This rank can be used to form a rank vector *R_jk_* such that:

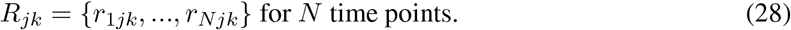

For each time series, a scalar *D_jk_* is calculated using the following formula:

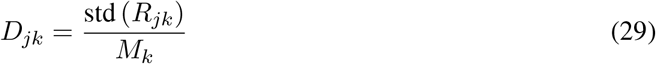

with std representing the calculation of a standard deviation and *M_k_* the number of time series data at a given treatment condition k. When two or more tumors at a given time point and treatment condition had the same recorded tumor volume, the average rank of these tumors was assigned for *r_ijk_*.

The outliers in the data are shown in Fig. 10 and were picked if they were singled out by the first criterion and/or second criterion. The exclusion rule for the second criterion was defined as *D_jk_* > 0.21.

## A.3 Additional figures

**Figure 11:**
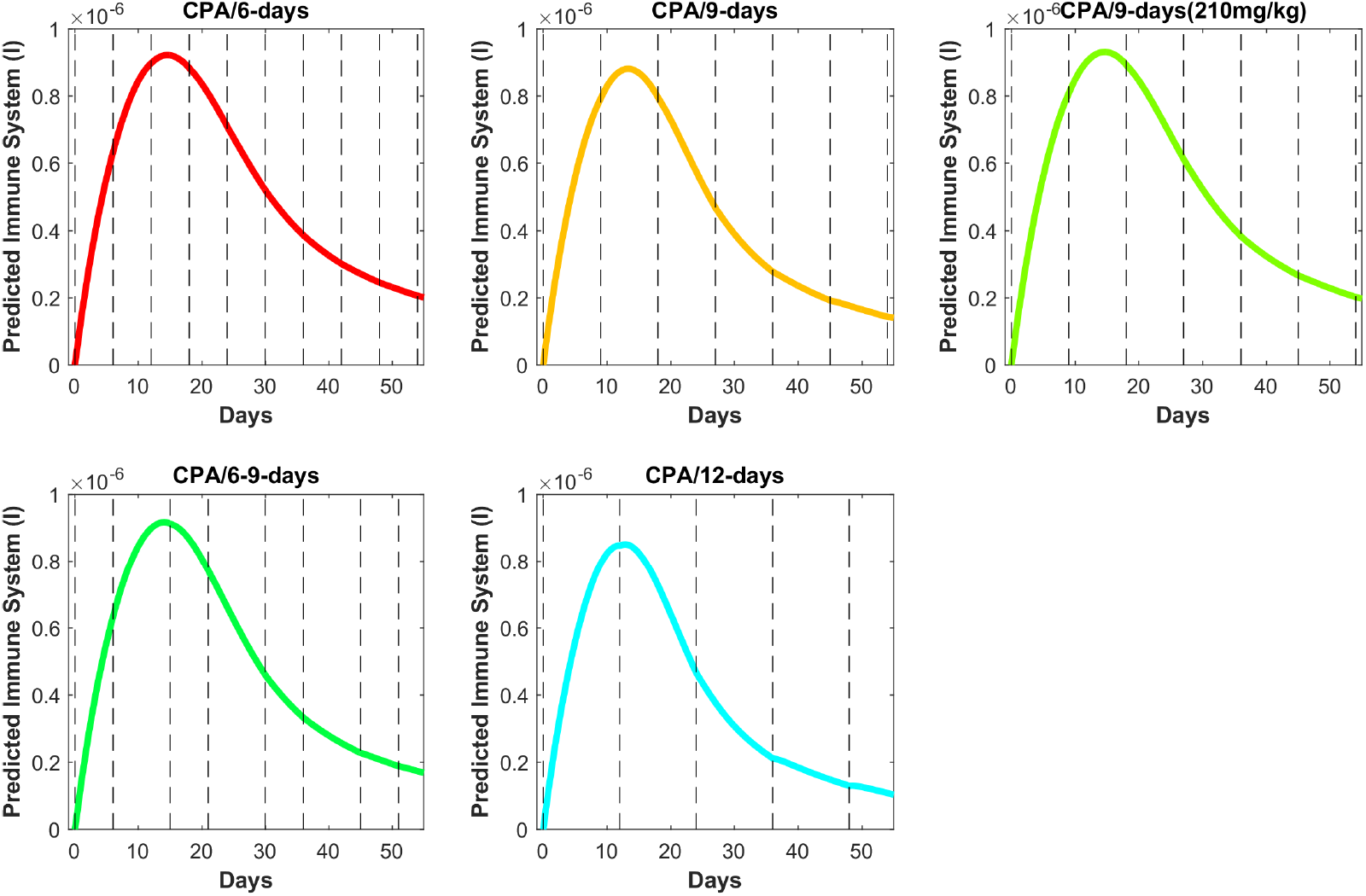
Predicted immune system from the model fits for the treatment conditions other than 1-CPA, 2-CPA, and 3-CPA.

**Figure 12:**
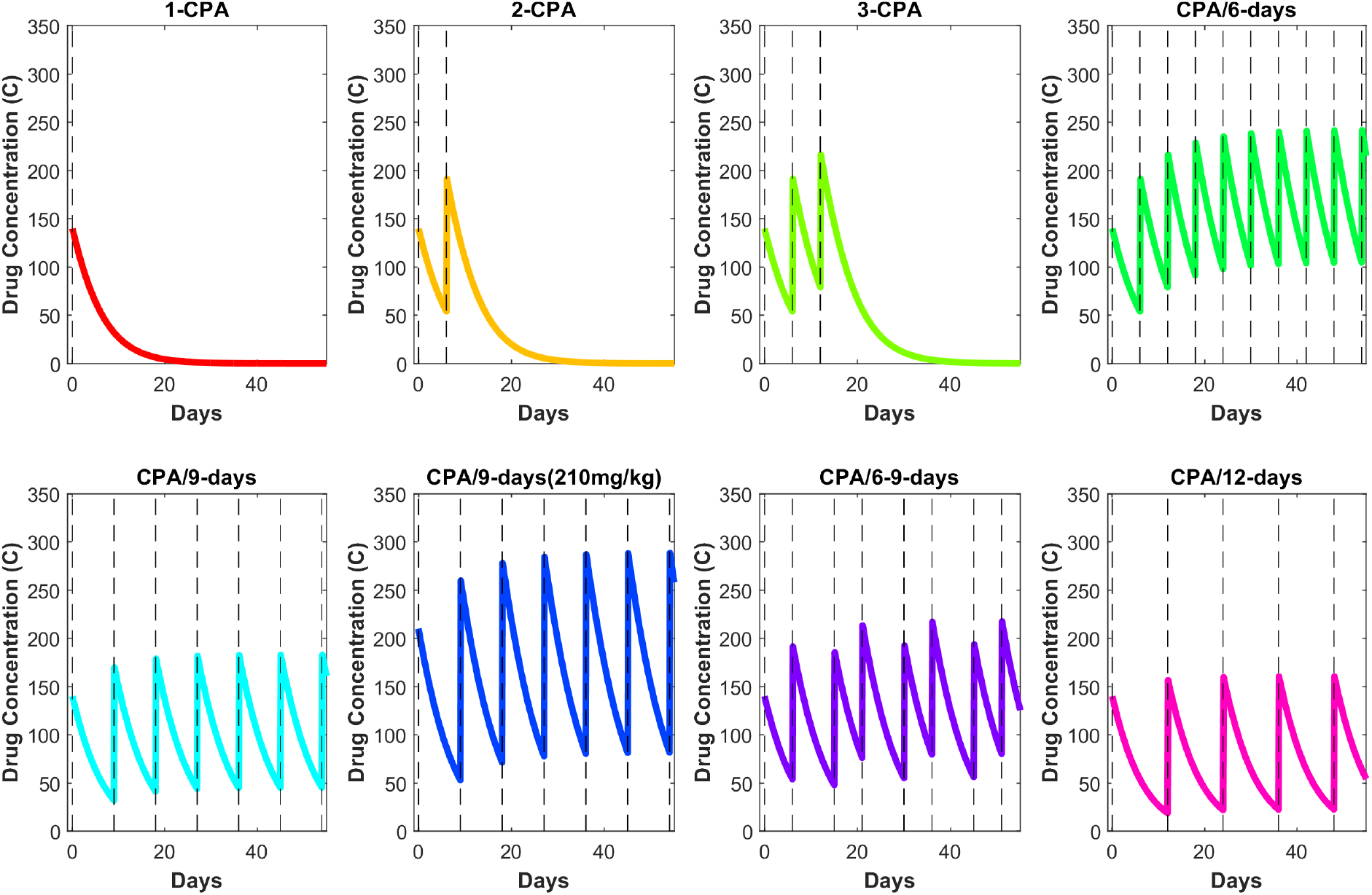
Prediction of the drug concentration (C) for all the experimental conditions in [9]

